# Detecting and quantifying overparametrization in RNA language models with REDIAL

**DOI:** 10.64898/2026.05.11.724344

**Authors:** Da Teng, Yunrui Qiu, Gokulakannan Sakthivel, Akashnathan Aranganathan, Lukas Herron, Pratyush Tiwary

## Abstract

While RNA language models (LMs) have served as foundation models (FMs) to advanced structural prediction, their evaluation relies heavily on supervised downstream tasks. Such tasks can often mask FM inefficiencies and reflect downstream training set memorization. To address this, here we introduce REDIAL (RNA Embedding perturbation Diagnostics for Language models), a zero-shot, unsupervised framework designed to extract coevolutionary signals directly from the high-dimensional latent spaces of RNA language models. By applying REDIAL, we uncover stark, layer-wise disparities in how popular RNA language models (LMs) internalize structural constraints through a layer-wise dissection and ablation study. Our results showed how such layerwise behavior deviates from protein LMs and is related to design flaws in the architectures. Specifically, we show that current RNA LMs are severely overparameterized relative to the limited sequence diversity of available RNA databases, leading to profound parameter inefficiency and overfitting. Furthermore, we establish that structure-guided pretraining fundamentally improves the signal-to-noise ratio of learned coevolutionary couplings compared to sequence-only baselines. Ultimately, this unsupervised evaluation paradigm exposes critical flaws in current parameter scaling strategies and provides a rigorous diagnostic benchmark to guide the development of more efficient, generalizable foundation models for RNA therapeutics and *de novo* design.

## Introduction

In the landscape of biomolecular structure prediction, language models (LMs) have emerged as powerful alternatives to traditional architectures, even as end-to-end generative models like AlphaFold 3^1^ achieve unprecedented accuracy. Language models are built on the principle that biological sequences can be encoded into high-dimensional latent representations through self-supervised pre-training^2^. These embeddings ideally distill the essential evolutionary and structural constraints of a sequence, serving as foundation models (FMs) that can be adapted for a wide variety of tasks. Leveraging this paradigm, ESMFold^3^ has achieved performance comparable to the original AlphaFold2 in protein structure prediction tasks, demonstrating that pretrained LMs can be useful to gain transferability from few-shot trainings. Following the success of ESMFold, numerous RNA language models have been trained^4–15^ as a promising framework for using limited training data to improve downstream predictions across diverse sequences. In terms of data availability, RNAs are different from proteins. Experimental RNA structural data and functional annotations are orders of magnitude scarcer than for proteins, but RNA genomic sequencing data remains abundant^16–19^.

Significant research effort has been directed toward understanding the success of language models in protein structure prediction. However, it remains an open question whether the same logic extends to RNAs. Zhang *et al*. recently demonstrated that protein language models (pLMs) implicitly encode evolutionary couplings (EC)^20^, providing a latent-space analog to the explicit multiple sequence alignment (MSA) inputs utilized by AlphaFold2 and AlphaFold3. Structure prediction via coevolution^21^ operates on the premise that if two residues frequently covary across evolution, they are likely to be spatially proximal. In proteins, this relationship is often anchored by a well-defined “native state” (or a discrete set of functional conformers), ensuring that coevolutionary signals reflect consistent physical contacts. While a subset of RNAs, such as ribozymes, riboswitches, and tRNAs, rely on specific tertiary structures to function^22^, many other non-coding RNAs lack a “native” fold. Instead, these RNAs transit rugged conformational landscapes characterized by multiple, competing alternative states^17, 18, 23–25^. While these RNAs may be promising drug targets, their structural heterogeneity requires a comprehensive characterization of their structure ensembles.^24, 26, 27^ This ensemble nature of RNAs structures may blur coevolutionary signals, and therefore poses distinct challenges for structure prediction^28^.

A second critical question regarding the development of foundation models is how to effectively assess their quality, either through explainable AI or through rigorous benchmarks. Like any deep neural network, the internal mechanisms of language models are often opaque; however, interpreting these mechanisms is essential for improving architectures and establishing reliability. Currently, state-of-the-art explanations of such Transformer architectures often involves using sparse autoencoders (SAEs) to separate the model activations into monosemantic features^29–32^. While SAEs allow for interpretation, they require significant training effort and possess inherent limitations. Once monosemantic features are extracted from activations, they must be correlated with functional annotations to be meaningfully explained. In large language models, this involves analyzing the specific contexts that trigger feature activation^30^, while in protein language models, it requires mapping these activations to known structural or functional motifs^31^. In any case, interpretation of these features ultimately requires sufficient high-fidelity structural or functional annotations, both of which are scarce for RNAs. Alternatively, some interpretations of these models use supervised models to connect self-attentions to contact maps, directly relating tertiary structures to learned evolutionary coupling. However, these interpreting supervised models, such as multilayer perceptrons (MLPs) or random forests, can be hard to explain themselves^33^. In terms of benchmarks, current language models are often assessed on downstream tasks, such as secondary structure prediction, tertiary structure prediction, or property prediction^6, 7, 34^. Because these supervised tasks require additional training data, these metrics fail to isolate whether performance gains stem from improvement of the language model’s core capabilities, superior downstream task optimization, or more concerningly, memorization of the downstream training data. Therefore, unsupervised evaluation frameworks must be developed to assess the intrinsic quality of pretrained language models, as they more accurately reflect the upper bound of information learned during pretraining. Such tests are highly valuable not only for benchmarking model performance and guiding the choice of model hyperparameters, but also for identifying potential directions for future improvements.

In this work, we address these two fundamental questions and provide new insights into current RNA language models. First, we propose an unsupervised REDIAL (RNA Embedding perturbation Diagnostics for Language models) algorithm to probe coevolutionary couplings. We then compare its performance on RNA LMs to the Categorical Jacobian (CJ) method developed for protein LMs^20^. We show that the proposed REDIAL approach has a greater signal-to-noise ratio compared to the CJ method. Next, using REDIAL, we highlight memorization issues in downstream structural prediction tasks commonly used to benchmark language models, and demonstrate how a purposefully designed training strategy can enhance model performance. Finally, we gain insight into what information RNA LMs learn and where it is located in the network by analyzing the layer-wise behavior of two RNA language models with REDIAL. The interpretation identifies abnormalities in current model architectures.

## Results

### Previous EC extraction methods are noisy when adapted to RNA LMs

Coevolutionary information is foundational to modern biomolecular structure prediction. While architectures like AlphaFold2 and AlphaFold3 utilize explicit MSAs as inputs to provide coevolutionary information, for recent LM-based prediction models such as ESMFold, it has been demonstrated that such information is implicitly encoded within their learned representations^20^. This emergent property arises because large-scale language models must learn the underlying geometry of the sequence space to produce meaningful embeddings, so as to achieve high accuracy in masked token prediction tasks. During training, the self-attention mechanism is forced to learn dependencies between tokens, including long-range ones that correspond to secondary and tertiary structures. This ideally provides the same coevolutionary information as performing explicit MSA searches in the genetic database used to train the language model.

The challenge, then, is how to extract this coevolutionary information from language models, ideally in an unsupervised way. Early works have found that specific attention heads of RNA language models can reflect underlying secondary structure^10, 35^. While such extraction is fully unsupervised, it suffers from high levels of noise. The likely reason is that although self-attention is an essential and interpretable mechanism for the language model to understand correlations between tokens in a sequence, the feed-forward network, which stores the majority of the information in the model^36^, is equally important. Therefore, a more holistic approach must be adopted.

In light of this, the “Categorical Jacobian” (CJ) approach (Figure 1) was developed for protein language models^20^. CJ resembles the class of linear surrogate models in the field of explainable AI research^37, 38^. The idea is to use a simple, yet interpretable linear model to approximate the local behavior of a complex model, utilizing the coefficients of the linear surrogate to derive mechanistic insights. By systematically perturbing different dimensions of the inputs and observing the corresponding output responses, one can determine feature importance: the larger the response, the more important the feature. For protein or RNA language models, where the inputs are biomolecular sequences, this approach translates to observing how the model’s output reacts to *in silico* mutations. Specifically, the CJ algorithm evaluates how the *logits* from the LM-head respond to mutations in the sequence (Figure 1). However, while this logit-based interpretation works well for protein vocabularies |𝒜| = 20, it introduces significant noise when applied to RNA language models due to their much smaller vocabulary |𝒜| ≈ 4. This is quantified by low bimodality (less capable of separating two modes) and low Kullback-Leibler distance to a Gaussian seen in CJ method (Figure S1). This establishes that for RNA LMs, CJ cannot effectively distinguish signal from noise. A detailed discussion of this discrepancy is in the Supplementary Information.

**Figure 1.**
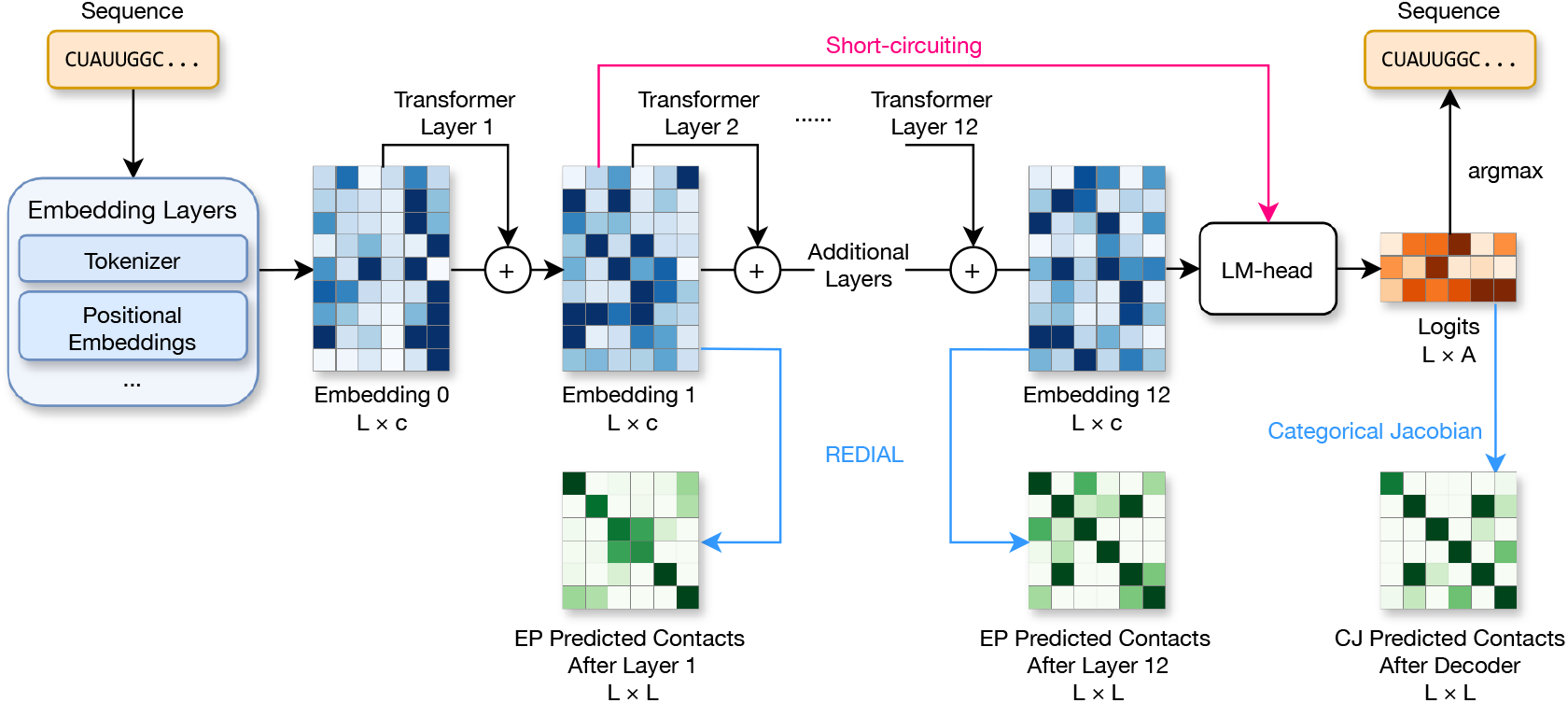
Typical architecture of a BERT-style RNA language model. Embedding Perturbations monitor the last layer of the model embeddings (and earlier layers for interpretations), while Categorical Jacobian monitors the logits output from the decoder. Short-circuiting is also used for layerwise interpretations later shown in this work. In the annotations *L* is the sequence length, *c* is the hidden dimension, and *A* is the number of alphabets.

### REDIAL: an unsupervised algorithm to extract coevolutionary signals from RNA language models

Since the perturbation in logits can suffer from noise for RNA alphabets, we look to find another representation produced by RNA LMs with high enough dimensions and thus noise tolerance. Therefore, we chose to monitor the sequence embeddings (aka hidden representations) produced by the model. The embedding layer and transformer stacks of ESM-2 (similarly illustrated in Figure 1) converts a sequence of length *L* into an 2-D embedding with dimensions *L* × *c*, where *c* represents the number of channels (*c* = 1280 for ESM-2-650M). This large number of channels ensures meaningful coupling signals will not be buried in model noise.

We first motivate this approach using the protein language model ESM-2-650M^3^ and Streptococcal Protein G, a well-studied system in investigating protein folding mechanisms. F30 is one of the hydrophobic cores important in the nucleation-condensation mechanism^39, 40^. If we look at the perturbation of the embeddings when we mutate the wildtype (WT) sequence to F30V (Fig 2a), clear signal bands are visible at the *i* ± 4 positions relative to F30. These localized perturbations reflect the *i* to *i* + 4 hydrogen-bonding pattern characteristic of an *α*-helix, suggesting that the model has internalized local secondary structural motifs. Notably, a distinct perturbation signal also appears at L5 (indicated by the red arrow). While distal in sequence, L5 is spatially proximal to Phe-30 in the tertiary fold, and also served as a hydrophobic core important in its folding^39^. This detection of long-range tertiary contacts indicates that ESM-2 learns residue-wise couplings from the massive sequence diversity in genomic databases, likely by capturing evolutionary couplings.

**Figure 2.**
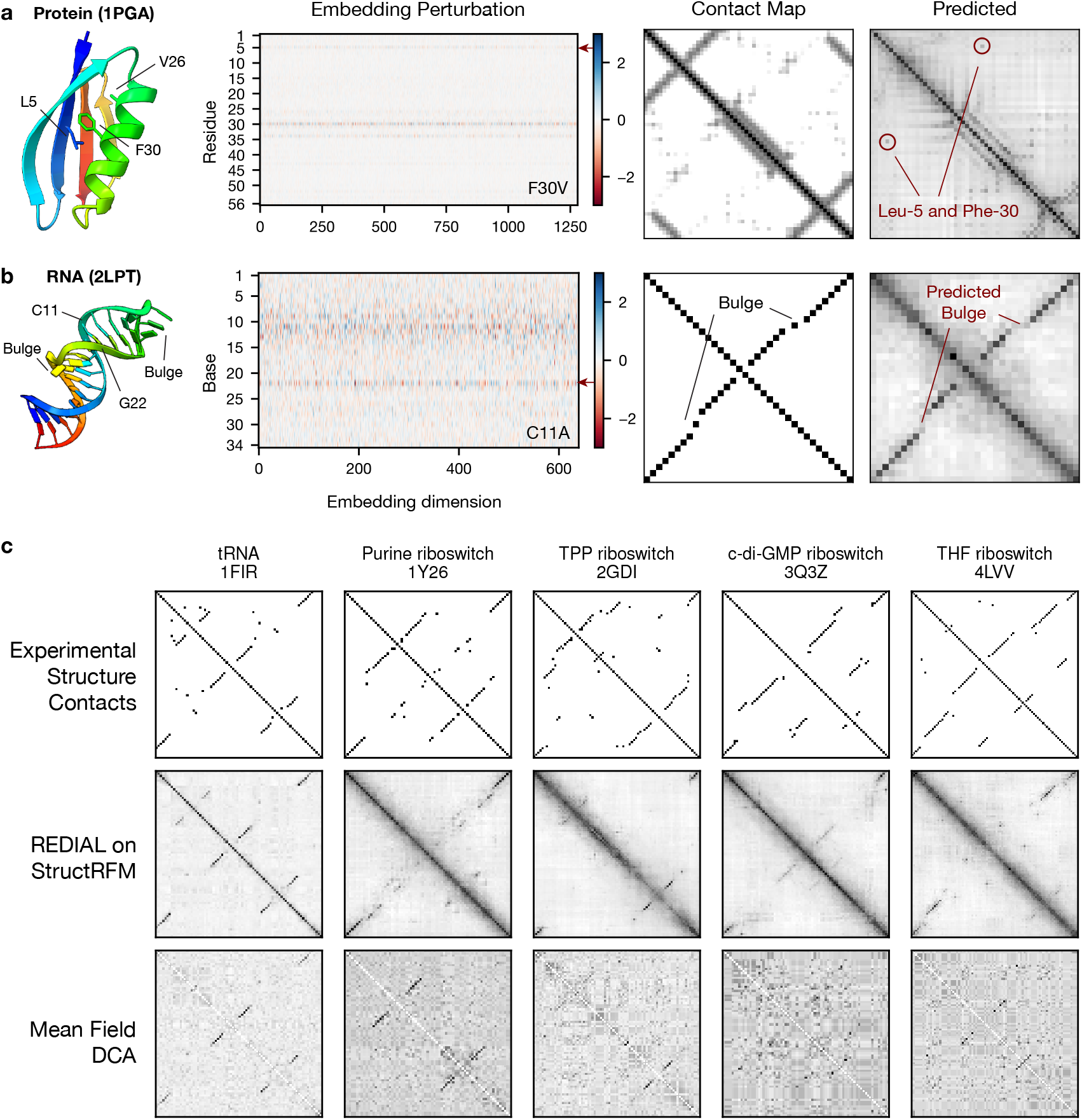
REDIAL algorithm. The REDIAL algorithm is first demonstrated on a protein, and then an RNA. (a) For the protein, F30 has secondary contacts (V26) and tertiary contacts (L5, red arrow). When mutating F30 to V, the embedding for both contacts see clear perturbations signified by a horizontal strip. (b) For the RNA, when mutating C11 to A, a strip of signal also shows up at G22, which forms a Watson-Crick pair with C11. These information can be compiled together to draw a coupling map (right, Predicted) that resembles the contact map, showing that secondary structures (*α*-helices and RNA bulges) and distal tertiary contacts can be recovered from REDIAL. (c) Validation of REDIAL algorithm in five systems shown in Weinreb *et al*.^22^ to have validated evolutionary coupling. We computed the contacts from experimental structure, the results from REDIAL, as well as a direct coupling analysis (DCA) from MSA alignments.

Motivated by these observations, we propose the RNA Embedding perturbation Diagnostics for Language models (REDIAL) algorithm. Specifically, for RNA, the alphabet 𝒜 consists of *A* = 4 standard nucleotides. For each of the *L* × *A* possible mutations (including the WT identity), we compute the perturbation in the encoded embedding, which retains the same shape *L* × *c* as the original embedding. This forms a perturbation tensor with dimensions [*L, A, L, c*]. The Frobenius norm is then calculated along the alphabet and channel dimensions, yielding an *L* × *L* coupling map. This matrix is subsequently symmetrized and subjected to Average Product Correction (APC, Equation 1) to produce the final coevolutionary coupling map^41^. A detailed mathematical description of the REDIAL method is provided in the Methods section. We emphasize that the simple nature of REDIAL means that it can be applied as a post-processing algorithm on generic language models for proteins, RNA and other modalities.

Here, to evaluate if the same algorithm generalizes to RNAs, we validated it using StructRFM^34^ on a 34-mer hairpin-like RNA intron (Figure 2b). StructRFM is a recently developed RNA language model based on the same BERT-style architecture as ESM-2, and was developed on the architecture of ERNIE-RNA^10^. It has a embedding dimension *c* = 640 and 12 layers of transformer, and we took the last layer of embedding to compute perturbation. Interestingly, this RNA contains two internal bulges, which induce a two-base register shift in the contact map relative to the diagonal. This shift is captured by our algorithm: when C11 is mutated to A on one arm of the hairpin, a significant perturbation is observed at G22, the shifted pairing partner, rather than the diagonal position 24. This demonstrates that REDIAL can extract non-canonical secondary structure features, such as bulges, from RNA LMs. By aggregating these individual mutational sensitivities, the REDIAL method generates a comprehensive coupling map that faithfully reflects the underlying secondary structure compared to the experimental contact map.

To further assess the consistency of this extracted coupling map with coevolution couplings from MSAs and contact maps from experimental structures, we applied the same method to five RNA systems previously characterized by Weinreb *et al*. to have verified, long-range coevolutionary couplings that are detectable via Potts models.^22^ We verified this with direct coupling analysis (DCA) to fit a Potts Model for these sequences using an MSA from RNAcentral where StructRFM was trained on, and the results were shown in Figure 2c. Due to the availability of MSA sequences, some predictions (3Q3Z, 4LVV) are noisy, but this should be treated as the upper bound of the coevolutionary information that could be possibly learned by any LMs trained on this dataset. These evolutionary couplings are mostly recovered by REDIAL algorithm (middle row, Figure 2c), but they still do not compare with the experimental contacts in the first row of Figure 2c.

### Memorization in downstream tasks necessitate unsupervised benchmarks

In the previous section, we demonstrated that while REDIAL-derived signals largely align with coevolutionary signals from DCA, they still fail to capture numerous contacts observed in experimental structures. This raises a perplexing question: how is the missing structural information inferred when a downstream model leverages a pre-trained foundation model (FM) for 3D structure prediction? For instance, RhoFold+ represents one of the earliest structure prediction models built upon a foundation model (RNA-FM), and it can incorporate supplementary data, such as an optional MSA, to generate 3D structures^6^. We seek to understand how information from these diverse inputs, alongside the internal weights of the downstream model, is synthesized during structure prediction tasks. Specifically, in cases of successful prediction, it is vital to discern whether the foundation model and MSA provide meaningful structural guidance, or if the model’s performance relies predominantly on the parameters in the structure prediction model.

To investigate this, we examine an instance of apparent memorization within the RhoFold+ pipeline. We evaluated this conjecture using the RNA-FM/RhoFold+ protocol on two well-characterized and deliberately selected systems: the HIV-1 TAR RNA and an mRNA AU-rich element (ARE). For the first system, the HIV-1 TAR RNA, numerous experimentally determined structures are available in the Protein Data Bank (PDB)^42–48^. However, the availability of homologous sequences is effectively zero; the default MSA generation protocol in RhoFold+ retrieved only 19 sequences, all of which are exact substrings of the original query sequence. This means no variability and coevolutionary information is available in the MSA. Consequently, the resulting MSA lacks sequence variability and provides no usable coevolutionary information. This limitation is reflected in our predicted coevolution map (Figure 3a) derived from the REDIAL algorithm and RNA-FM, which confirms that the pretrained language model did not capture any evolutionary couplings for this target. Strikingly, when the downstream RhoFold+ model was used to predict the 3D structure, both with and without the sparse 19-sequence MSA, the resulting predictions closely matched the experimental structure. This indicates that the downstream module successfully recovered the conformation without relying on coevolutionary signals.

**Figure 3.**
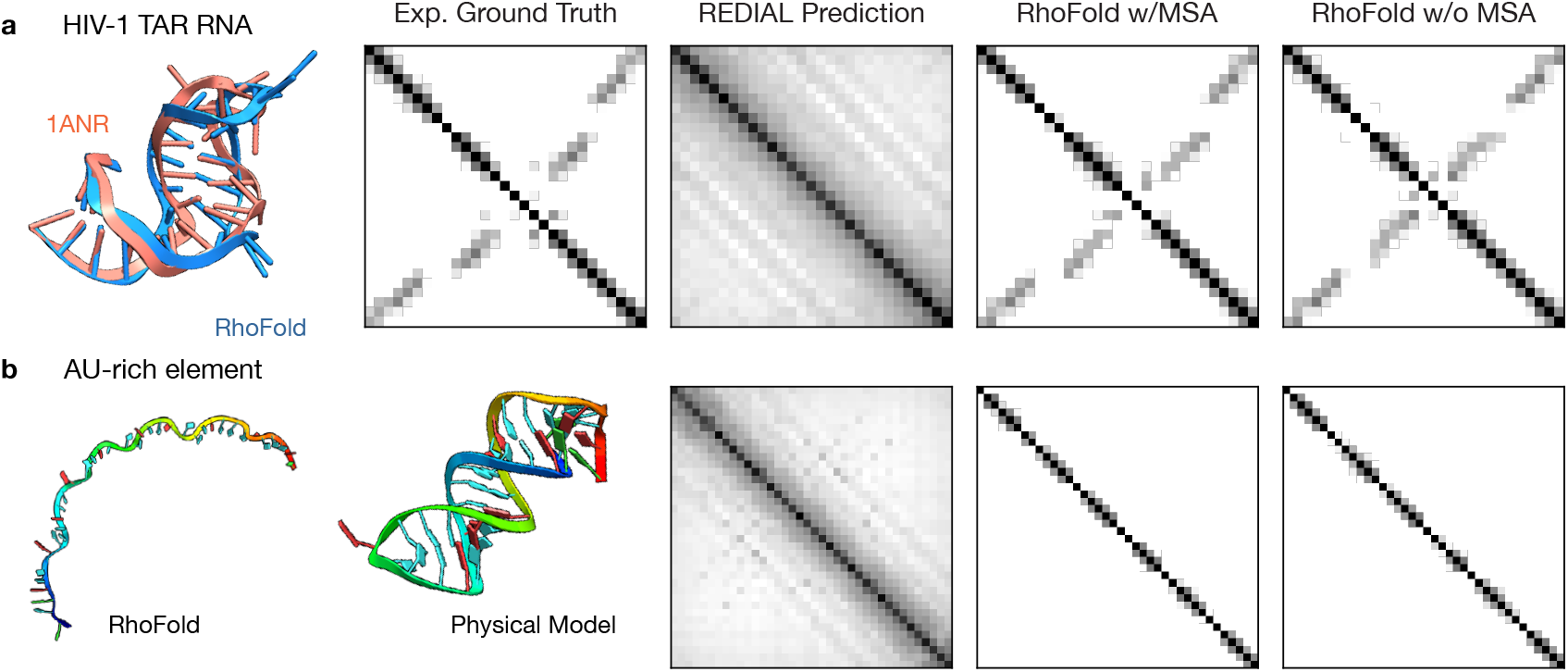
Downstream structure module memorizes structures without learning physical constraints. For two systems, we compared the structure predicted by RhoFold+, a downstream structural prediction model for RNA-FM, with either experimental structure or the physically predicted structure with thermodynamics scores. In both cases, REDIAL suggests no EC signals were learned, and thus passed down to the downstream modules. (a) For HIV-1 TAR RNA, RhoFold+ recapitulated the experimental structure either with or without additional MSA. (b) For the AU-rich element of messenger RNAs, RhoFold+ predicted a linear polymer in either cases. All contacts shown here are the distograms of the 3D structures, where a cutoff to white is set at 8Å.

We then investigated whether RhoFold+ relies on canonical base-pairing rules and physical principles to make such a successful prediction, or if it simply memorizes training data. To test this, we evaluated RhoFold+ on the ARE, comparing against published results by Wilson and co-workers^49, 50^ (further system details in Methods). Since RNA-FM was trained exclusively on non-coding RNAs and no experimental structure for this sequence exists in the PDB, it was likely unseen by both the language model and the downstream folding model during training. This is first proved by an MSA search with RhoFold+, yielding zero homology sequence. In Figure 3b, REDIAL confirmed again that no coevolutionary signals could be extracted from the language model. Different from the HIV-1 TAR case, RhoFold+ predicted this RNA to be a linear polymer without any secondary structure. This result not only contradicts the fundamental biochemistry principle that Adenine and Uracil can form Watson-Crick base pairs, but is also inconsistent with previous experimental folding evidences: Wilson and co-workers previously employed mFold^51^ to predict the minimum free energy secondary structure and obtained a folded conformation^50^ that is highly consistent with previously reported nuclease-mapping experiments^49^. This predicted secondary structure further provides a template for subsequent tertiary-structure reconstruction using FARFAR2^52^, yielding a compact, well-folded conformation (Figure 3b, second column; see Methods for additional details on structure prediction), which directly contradict the result from RhoFold+.

These two RNAs were carefully chosen to serve as representative systems. HIV-1 TAR RNA is evolutionarily out-of-distribution (in the genetic database), but structurally over-represented in the downstream task’s training set. The AU-rich element is both evolutionarily and structurally out-of-distribution, but has obvious secondary structure potentials that can be easily captured by physics-based models. This isolates the information available to this downstream structure model so we can confirm that the correct prediction for HIV-1 TAR is from memorization. Many foundation models are now benchmarked on downstream tasks (e.g. BEACON^53^, RNA-Puzzles^54^), which can obscure whether high performance stems from generalized learning or mere memorization. While these observations are not intended to suggest that memorization is the universal operational mode of these models, the existence of this discrepancy highlights the need for benchmarks that rigorously evaluate the quality of the foundation model itself, rather than the downstream task. In this context, our REDIAL algorithm offers a robust measure of the learned coevolutionary information of the language model.

### Structure-guided language model learns better basepair couplings

Given that most current RNA LMs, such as the aforementioned RNA-FM and StructRFM, are already built upon state-of-the-art RNA sequence databases^55^ and the proven BERT architecture, a central question in the development of RNA LMs is how to drive further optimization. Unlike LLMs in the natural language domain, the available biological sequences as training data are finite and cannot scale as rapidly. In terms of architectures, debate continues regarding the relative merits of masked language models (MLMs) versus decoder-only causal architectures^56, 57^, both paradigms have been extensively explored in the context of biomolecular sequences. Consequently, the next significant performance gains may instead reside in the refinement of the training strategy.

For instance, the advance of StructRFM^34^ is to explicitly integrate predicted secondary structural priors into its pre-training phase. Rather than relying solely on stochastic token masking, the model employs a structure-guided masked language model (SgMLM) strategy. If a randomly selected base is involved in a known base pair, its corresponding structural partner is also masked. This approach forces the model to infer tokens not just from canonical pairing rules, but from hidden coevolutionary couplings. Other than the training strategy, these two models remain largely the same, especially in architecture. They have the same 12 layers of self-attention, while StructRFM has slightly higher hidden embedding dimensions. StructRFM reported significant improvements in downstream structural benchmarks^34^. However, it remains unclear if these gains reflect a more robust foundation model or better task-specific fine-tuning.

By applying REDIAL to both RNA-FM and StructRFM, we aim to compare the coevolutionary signals learned by these two models. This allows us to decouple the representational quality of the foundation model from the performance of downstream predictors, providing a direct measure of how structure-informed training influences the internal understanding of RNA informatics. To compare REDIAL predicted coupling with ground truth contact maps computed from experimental structures, we must define a metric that evaluates how a continuous score reflects a binary contact map. There are two primary challenges in defining this metric: First, the range of coupling strength varies between systems and language models, making it impossible to determine a single cutoff value to binarize the predicted couplings for comparison. Second, picking a fixed percentage of the highest predicted couplings is infeasible, as the percentage of native contacts depends on the length of the RNA (Figure S6). To address these issues, we adopted the Area Under the Precision-Recall Curve (PR-AUC) as our metric. This approach evaluates the utility of a continuous score in a binary classification task while accounting for class imbalance, where 0 is the expect score of a random prediction, and a score of 1.0 means perfect prediction. More details are described in the Methods section.

We compared how RNA-FM and StructRFM perform on our NAKB test set consisting of 533 structure-sequence pairs (see Methods for data curation process). We found that by incorporating structural data, the resulting scores significantly improve in StructRFM compared with the sequence-only model RNA-FM (Figure 4a, S3, S4, S5), almost every point shows a higher score for StructRFM than for RNA-FM. These improvements can be categorized into two types. The points clustered near the *y*-axis represent cases where RNA-FM fails to predict meaningful contacts, but StructRFM succeeds. An example of this scenario is shown in Figure 4b (`1EBS`), where RNA-FM cannot identify any evolutionary coupling, but StructRFM correctly predicts the primary stem (the diagonal) and the bulge-induced shift in the diagonal contacts. The points closer to the dashed diagonal line (gray, Figure 4a) are systems where both models perform well, but StructRFM still yields a measurable improvement (Figure S4). An example of this is the tRNA 6UGG in Figure 4c, where the predicted couplings from StructRFM score higher due to a better signal-to-noise ratio. It can be seen from the two predicted contact maps that StructRFM produces much cleaner, sharper predictions, such that the precision-recall curve of RNA-FM starts to drop at smaller recall percentages.

**Figure 4.**
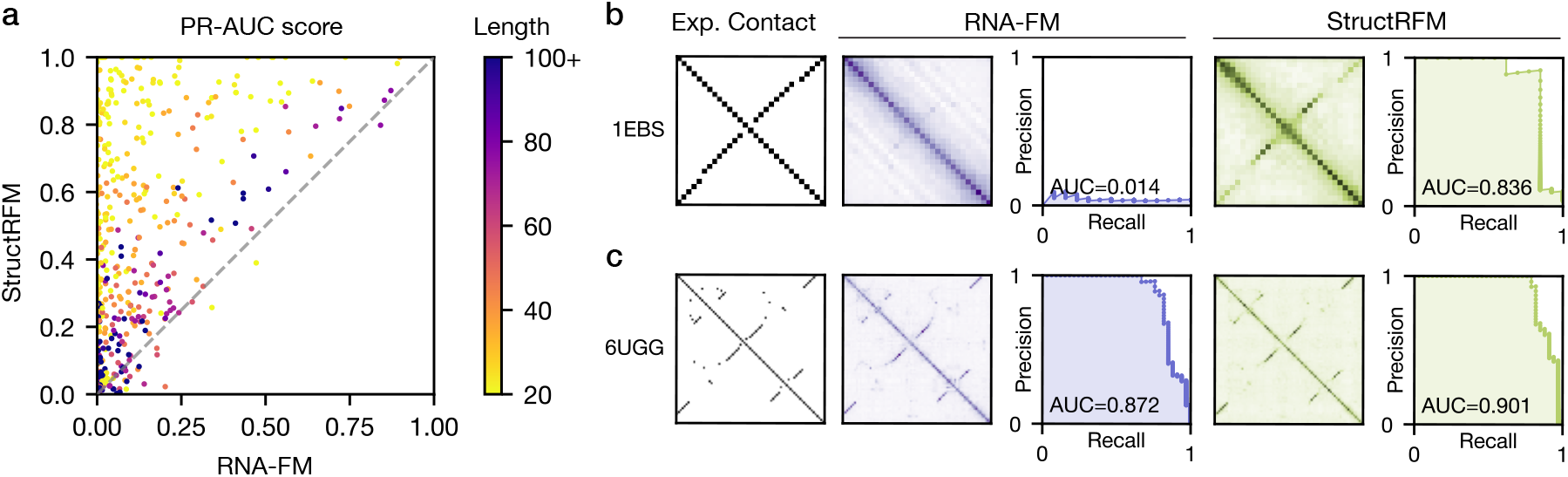
Structure-guided model outperforms RNA-FM in contact prediction comparing with experimental structures. (a) Comparison of the PR-AUC score between structure-guided StructRFM and RNA-FM, which is trained on sequences only. The higher the score, the better the REDIAL predicted contact recapitulates the experimental contacts. Each dot is one entry in our curated NAKB subset, with color denoting its sequence length. (b) For a hairpin, StructRFM predicts the secondary structure (including the bulges) that RNA-FM cannot predict. For RNA-FM, the precision consistently close to a random number generator. (c) For a more complex tRNA, both models predict the correct contacts, but StructRFM predictions are higher in signal-to-noise ratio. This gives higher precision at the same recall level, resulting in a higher area-under-curve (AUC).

A particularly interesting observation from Figure 4a is that the systems where the sequence-only RNA-FM fails badly are those with shorter sequence lengths (yellow dots) rather than longer sequences (dark blue) (more in Figure S5). This shows that sequence context alone is not enough to reliably extract coevolutionary signals from short sequences, while structure-guided training can overcome this limitation. This trend is reflected in protein-related tasks as well, where ESMFold performs much worse on short peptides than on longer, structured proteins^58^. For short sequences, the chance of two non-homologous sequences that follow very different evolutionary paths being spuriously associated is much higher. In other words, the mutual information is more likely to be buried in the high entropy of the sequences. Such noise can be more prominent in RNAs than in proteins, because the vocabulary size of RNAs (*A* = 4) is much smaller than that of proteins (*A* = 20). That said, most long sequences, especially those with *L >* 60, still do not get good prediction from either models.

### Layerwise interpretation of RNA LMs

Another benefit of our REDIAL algorithm is the ability to look at how each self-attention layer of the encoder behaves, and potentially analyze how information was picked up through the 12 Transformer layers. Figure 1 shows a typical architecture of a BERT-style language model. In such architectures, each Transformer layer updates the hidden representations (embeddings) iteratively, so these intermediate embeddings can be extracted for analysis. In total we performed two analysis. First, we “truncate” the model by using REDIAL on intermediate embeddings, and calculate the PR-AUC score for the resulting coevolutionary maps (Figure 1, blue arrows). This will give us insight on how much each layer contribute to the final coevolutionary map at the last layer. Second, we “short circuit” some of the layers and feed those representations directly to the original model decoder to recover the logits. The increase in the model perplexity, which is defined as a measure of the model’s uncertainty in predicting the true sequence, can tell whether we lost important information from the embeddings. These two tests correspond to different learning objectives. The scores from recovered coevolution test whether the model learns relationships between bases, while the perplexity test tells whether each layer is needed for recovering the sequence.

We tested these two operations on both StructRFM and RNA-FM and our NAKB test set, and we found that despite having both 12 Transformer layers in their encoder, these two models were trained to learn the coevolutionary in very distinct ways. In the first test (Figure 1a), we found that StructRFM is “tail-heavy.” It picks up negligible information in the first ten layers and most of the information about coevolution is gained in the last two layers. In contrast, RNA-FM is “head-heavy.” It learns most of the coevolutionary information in the first six layers, after which the scores barely improve further. In light of this observation, we short circuited the less performing layers in each model in the second test (Figure 1b). First *x* layers are short circuited from the tail-heavy StructRFM and its performance does not decrease significantly until *x* = 7. In RNA-FM, where the last *x* layers are less informative, no significant reduction in information is observed until *x* = 4.

To understand these behaviors better, we took 4 examples from StructRFM and showed them in Figure 5c. The system `1AJF` has a simple hairpin secondary structure, and `4PLX` is a complicated triple helix. Whether the model has learned the correct contact (hairpin) or not (triple helix), no useful information emerged until after Layer 11. The case is different for tRNAs, where information starts emerging very early for `1FIR` and *6UGG*. As early as Layer 2, blocks of black shades have appeared corresponding to different segments of the secondary structure. Such information is refined and “denoised” over the later layers. Layer 9 in StructRFM seems to do more damage than help, as shown in Figure 5b that ablating the first nine layers results in a better model than ablating the first eight. This can also be seen in the case of `1AJF` and `4PLX` in Figure 5c that Layer 9 dramatically increased the noise near the diagonal, and Layer 10 to 12 tried to eliminate it but cannot eradicate it.

**Figure 5.**
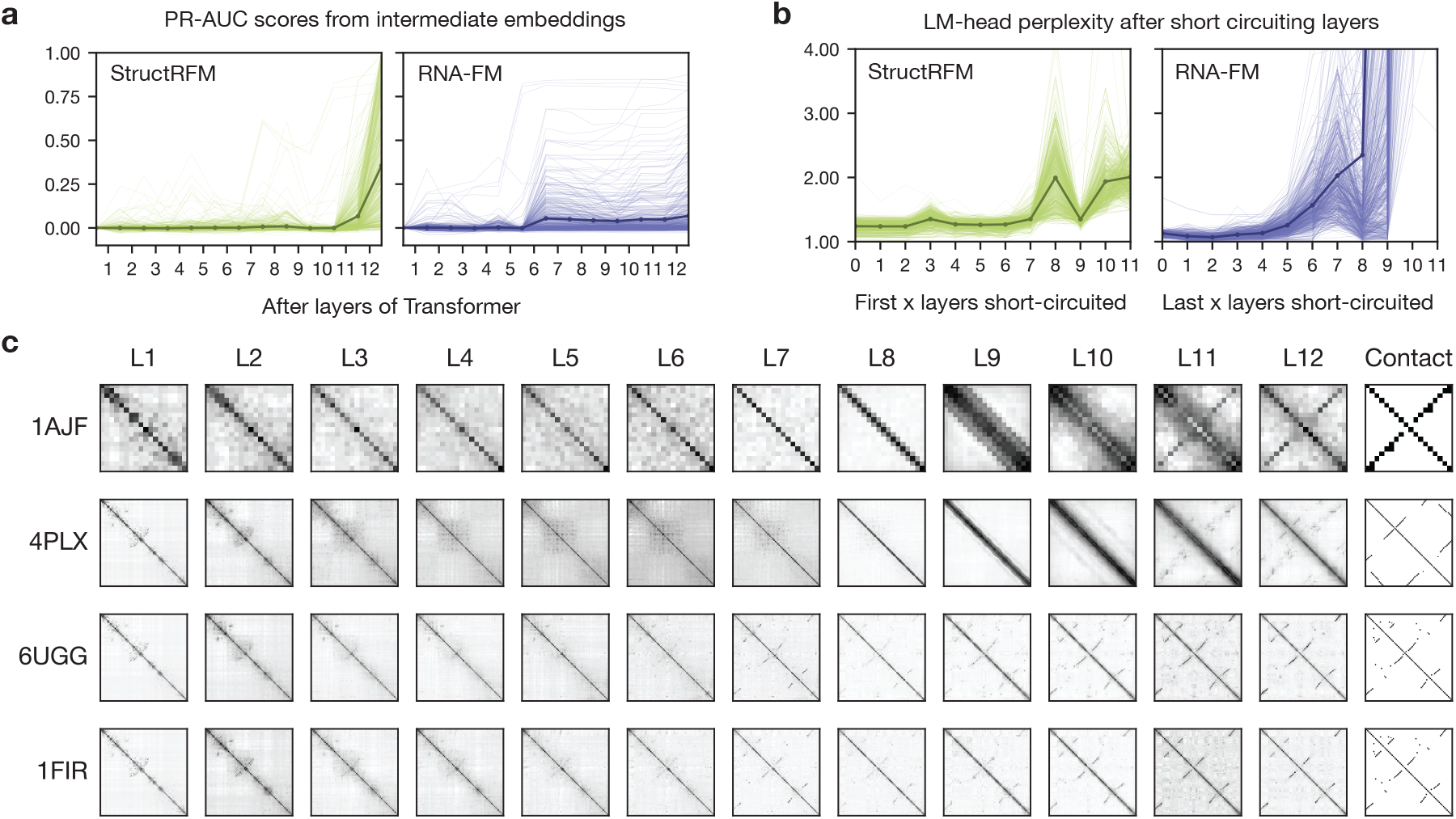
Layer-wise interpretation of RNA language models reveals functional imbalances across layers. Analysis of intermediate representations using the curated NAKB subsets. (a) PR-AUC scores derived from intermediate layer embeddings. StructRFM captures minimal coevolutionary signals until Layer 11, whereas RNA-FM performance plateaus after Layer 6. (b) LM-head perplexity following the short-circuiting of Transformer layers. For StructRFM, short-circuiting the first 6 layers has minimal impact, while altering the latter 6 layers yields mixed effects on the LM-head. For RNA-FM, short-circuiting the final 8 layers incrementally increases perplexity, while the first 4 layers are more essential. In both (a) and (b), thin lines represent individual RNA systems, and the bold line with markers denotes the mean across all systems. (c) Representative examples of contact maps extracted after each StructRFM layer. The final column shows the ground-truth contact map derived from the PDB. For targets 1AJF and 4PLX, structural information is predominantly acquired in the final layers (L11 and L12). Conversely, targets 6UGG and 1FIR are two examples that accumulate this information more gradually across the network depth.

### Evidence of overparametrization in current RNA LMs

One might expect that well-trained language models follow a process of iterative refinement, gradually enriching the residual stream with semantic information through successive self-attention layers. However, previous studies on language models for either natural language^59–61^, protein sequences^31, 62^, and RNA sequences^10^ have demonstrated this is not the case. Depending on the specific architecture, most of the functional information usually resides within either the middle layers (for autoregressive models) or the late layers (for BERT).

In contrast, our results in the preceding section revealed that the two models under investigation displayed highly non-standard and convoluted behaviors. First, coevolutionary information does not accumulate incrementally. Despite StructRFM displays a tail-heavy learning as expected in BERT-style language model (Figures 5a, b), specific layers appear to capture spurious or incorrect information. Second, we observed that RNA-FM, contrary to the typical late-layer dominance of BERT-style architectures, exhibits atypical “head-heavy” characteristics where the early layers perform the vast majority of the functional computation. We will further investigate these anomalous behaviors in this section.

First, since StructRFM is tail-heavy, we constructed shortened models by removing the first six layers and various combinations of the late layers in StructRFM. For these “short” models, we evaluated their perplexity and contact recovery capability. We systematically experimented with removing any possible combinations of the last 6 layers. As expected for a tail-heavy architecture, we found that removing the first 6 layers alone (model L7-L12) has minimal effect on these two metrics (Figure 6a, b). However, further reducing the model to five layers starts to significantly compromise its performance, mostly in its capability to predict evolutionary coupling. The best three five-layer models in contact scores all contain Layers 8, 9, and 12, highlighting their potential importance. This observation agrees with the calculated Shapley values for each layer (Table 1). Layer 8, 9, and 12 contributed the most to the contact score, despite we observed in Figure 5a that noisy signals were picked up near the diagonal at Layer 9. The complicated matter comes from Layer 10, where it seems to be contributing less than the previous and succeeding layers, exhibiting a non-monotonous behavior. In terms of model perplexity, only the last two layers have contributed to reduce the model perplexity.

**Table 1.** Shapley values of StructRFM late layers. The Shapley values of each of the last six layers of StructRFM regarding their contributions to the model perplexity (lower the better) and contact scores (higher the better).

| Layer | Perplexity (↓) | Contact Score (↑) |
| --- | --- | --- |
| 7 | 1.6455 | 0.0299 |
| 8 | 0.7975 | 0.0568 |
| 9 | 2.0835 | 0.0718 |
| 10 | 0.1052 | 0.0428 |
| 11 | -0.3697 | 0.0513 |
| 12 | -2.9941 | 0.0582 |

**Figure 6.**
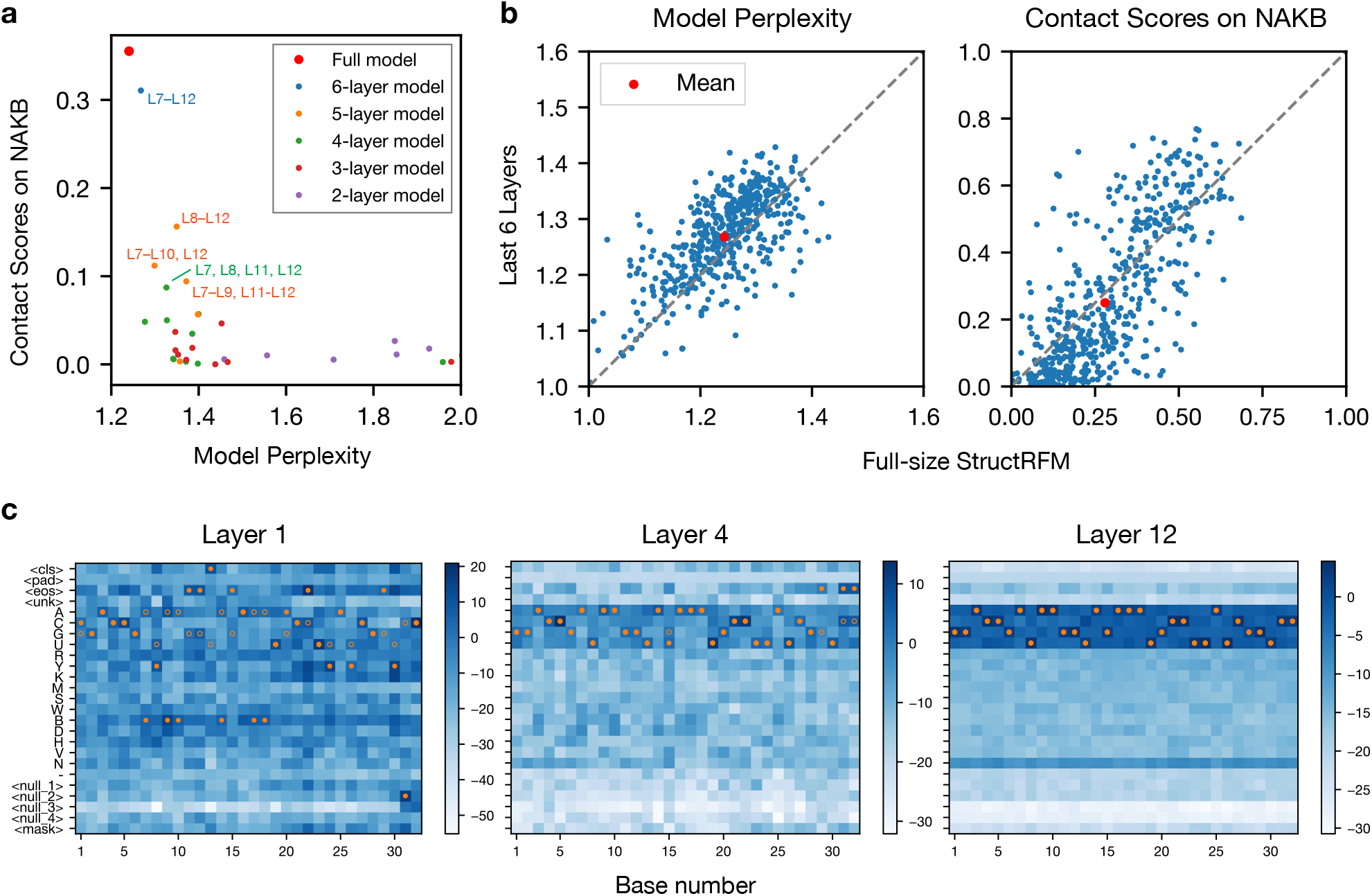
Ablation studies reveal the layer-wise mechanisms of RNA language models. (a) Systematic screening of truncated models derived from the final 6 layers of StructRFM. Models are evaluated based on their contact prediction capability (PR-AUC scores on REDIAL) and sequence recovery (model perplexity). (b) Performance comparison of the isolated last 6 layers versus the full-size StructRFM on the curated NAKB subset. Red markers denote the mean, demonstrating that the truncating half of the model does not deteriorate performance significantly. (c) Decoded logits from intermediate embeddings of RNA-FM. Hollow circles indicate the ground-truth base, while solid circles represent the model’s primary prediction (highest logit value). Overlapping circles denote a correct prediction. Background heatmaps display the logit values, with darker blue corresponding to higher values. The analyzed sequence is derived from PDB `2LI4`.

The second experiment investigates the uncommon head-heavy behavior of RNA-FM, as well as the exploding perplexity observed when ablating more than the first eight layers. We hypothesize that this is due to the different vocabulary sizes of the two language models. In RNA-FM, there are a total of 15 letters for nucleotides (compared with five in StructRFM), including many non-standard, low-frequency bases like H, V, *etc*., as well as four null tokens and a gap. Decoding the intermediate embeddings and examining the corresponding logits (Fig. 6c; Fig. S7) reveals a two-stage predict behavior. After a single Transformer layer, the model is very undecided on its predictions. In the early stage (i.e., Layer 1 to 4), the model has almost recovered the correct sequence, but with lower confidence, as the log-likelihoods for the correct base are not very outstanding. The later layers do not contribute much to identifying the correct base, but mostly sharpen the output distribution by concentrating probability on the most common tokens including ACGU and N. Consequently, ablating the later layers hurts the model very minimally, but if we ablate before Layer 4, more uncommon bases or even auxiliary tokens will be predicted, drastically increasing the model perplexity.

In summary, our layerwise analysis reveals that neither model utilizes its full depth to continuously refine structural and functional representations. StructRFM concentrates its contact prediction and sequence recovery capabilities into a few critical late layers, rendering its early representations largely redundant for these tasks. Conversely, RNA-FM demonstrates a head-heavy computation where early layers establish the sequence, and late layers seemingly act only as a probability filter against rare vocabulary tokens rather than extracting deeper coevolutionary insights. Together, these unique layer utilization patterns provide strong evidence of model overfitting and highlight significant parameter inefficiencies in current RNA language model architectures. Meanwhile, we realized that in the RNA language model developments, systematic benchmarks on the choices of model architectures have never been reported. These choices include the number of transformer layers, size of vocabulary, as well as issues we haven’t covered in this paper, such as size of feed-forward networks, or positional embedding types. From our analysis, we propose that a shallower transformer stack and smaller vocabulary with more training epoches may help the models with the same amount of training effort. We will detail this in the following discussion.

## Discussion

Since the introduction of transformers, many highly capable language models have been built for both human languages and macromolecular sequences^63^. The task of interpreting and understanding these massive architectures often intertwines with finding ways to improve them further^30, 31^. In this paper, we introduce REDIAL, an unsupervised, model-agnostic benchmark designed for this exact purpose. By extracting coevolutionary signals from the models, we can both assess model quality and interpret their underlying working mechanisms. Our contributions are threefold. First, we developed an unsupervised score that measures how much coevolutionary information a model has learned, which is an essential metric for structural prediction and a core reason these models are effective. Second, we used this tool to probe the layer-by-layer information flow of two RNA language models, StructRFM and RNA-FM. Finally, inspired by these findings, we conducted an ablation study highlighting critical issues regarding vocabulary and model sizes.

The REDIAL algorithm displayed pronounced capability in recovering details in the contact maps, and is the best unsupervised method known to us to predict contact maps from language models^10^. For instance we showed how it can recover the bulge-induced pairing shift in hairpin-like secondary structures (Figure 2b, 4b as examples). However, it is unknown if such information comes from intrinsic coevolution signals learned and amplified by the models, or the secondary structure information injected in the training of StructRFM, as there is a clear discrepancy between the performance of RNA-FM and StructRFM (Figure 4a), especially on shorter RNAs. Architecturally, RNA-FM included many infrequently used tokens and exhibits unexpected behavior as found in our layerwise analysis (Figure 6c). This choice, and other training hyperparameter choices, can result in differences in model capability to amplify existing coevolutionary signals.

Applying this benchmark yielded a crucial, alarming conclusion: evaluating language models purely by downstream tasks is dangerous due to severe risks of memorization. In our paper, we demonstrated this by analyzing RhoFold+, a downstream task accompanying RNA-FM, which memorizes its own training set (HIV-1 TAR RNA), and predicts a linear polymer for an obviously folded RNA, all without learning any underlying base-pairing rules. Such memorization can also plague tertiary structure prediction, sequence classification, and many other property prediction models. This underscores the necessity of our REDIAL algorithm and calls for a paradigm shift toward evaluating these models through an unsupervised lens.

Our analysis of model overfitting also provides crucial guidance for future model training. Two landmark papers set the stage for the scaling laws of large language models: the first one in 2020 from OpenAI posited that bigger models are inherently better^64^, while the second paper in 2022 from DeepMind^65^ searched for the optimal ratio of model size to available training tokens to maximize the utilization of data and compute. In the 2022 DeepMind work, they demonstrated that in the era of GPT-3, most large language models were significantly overparameterized relative to the amount of available training tokens. This phenomenon extends beyond natural language; for instance, an analysis by Zhang *et al*.^20^ explored the relationship between model size and the potential amount of coevolutionary data available in human genomic databases, arguing that the model size at which ESM-2’s performance starts to taper off (3 billion parameters) is on the same order of magnitude as the estimated coevolutionary contacts. Similarly, within RNA-related tasks, the authors of trRosettaRNA2 also confirmed a “minimal performance impact” when reducing the number of Transformer blocks from 48 in their predecessor version to 12 in the current version^66, 67^. Aligning these historical precedents with our own observations in the ablation and layerwise analyses of RNA-FM and StructRFM, we conclude that current RNA language models are likely suffering from a similar overparameterization problem to GPT-3-era large language models.

To be specific with numbers, the two models we tested have roughly the same number of parameters, with StructRFM containing 86M and RNA-FM containing 100M parameters. Due to differences in floating point precision, the raw bit size of RNA-FM’s weights (roughly 9.55 billion bits) is substantially larger than that of StructRFM (2.75 billion bits). Meanwhile, based on the original report of RNA-FM^6^, we estimate that the total entropy of its training set has an upper bound of 33.8 billion bits, assuming a random distribution of alphabets (with biological patterns in the sequences further decreasing this entropy). Even compared to the upper bound of the genetic database’s information content, the model size of RNA-FM is strikingly large. With a storage capacity capable of holding more than a quarter of the total dataset entropy, the network possesses enough bandwidth to simply memorize a vast portion of the training sequences.

Beyond raw storage constraints, we can also analyze this severe overparameterization from the perspective of effective information capacity. Current estimates for natural language LLMs suggest that models can memorize factual knowledge (that cannot be reproduced from reasoning within the model) at a rate of roughly 2 bits per parameter^68^ to 3.6 bits per parameter^69^, giving these models hundreds of millions of bits of effective memorization capacity. Applying this metric, the factual information capacity of these two RNA LMs is estimated to be ∼300 million bits. However, estimating the information content of RNA contacts following the discussion in Zhang *et al*.^20^ yields a required capacity of a different order of magnitude. Assuming 4,300 RNA families (approximating the number in the current Rfam database^70^), an average length of 64 bases, and two contacts per base, we calculate a target capacity of a mere 8.8 million bits of information. This is roughly the relational information required to learn coevolution, which is strikingly small compared to the capacity of the model. Transformer feed-forward networks (FFNs), which dominate the parameter count, function primarily as extensive key-value memory banks^36^. Because language models tend to fill their available information capacity with memorized data before generalized learning (or “grokking”) occurs^69^, the vast excess capacity in the FFNs provides a potential shortcut to “hack” the masked language model objective. Rather than forcing the self-attention mechanisms to learn fundamental evolutionary couplings, the model may be using the FFNs to memorize sequences and work on pattern matching. This also provides some clues on why attempts to solely compare attention maps to contact maps do not yield results as good as we get in this paper.^10, 33^

With all that said, unlike the natural language field where a company can enrich their training data with a high-speed scanner and millions of unconsenting books^71^, RNA sequences remain stubbornly expensive and slow to source in bulk, so it is unlikely we will see the same convenient, exponential leaps in available genomic sequences. Therefore, the only practical way to train better models is through regularization in network architectures or better training strategies. We have already seen StructRFM incorporating secondary structure information into their training strategy. Their structure-guided masking strategy essentially augmented the relatively scarce training data, injecting distilled information from another model or physical calculations into the language model. In our work, we have shown their model outperforms the naively trained RNA-FM by a significant margin, especially on short sequences where coevolution from mutual information is noisy.

Lastly, it is important to acknowledge the practical reality that current RNA language models are mostly built in academic institutions with limited computation power and engineering effort, unlike the massive, industry-backed protein language models like ESM. As the RNA community cannot simply rely on brute-force parameter scaling, the path forward requires a highly innovative approach in architectural design and training strategy to build robust, reliable tools that the scientific community can confidently rely on.

## Methods

### 1 Curation of the dataset from NAKB

We sourced our RNA structural sets on Nucleic Acid Knowledgebase (NAKB)^72^ on February 2, 2026. We looked for all systems with single chain without other modalities such as protein or small molecules. We also removed structural duplicates by searching inside the “Representative RNA Set” of NAKB. This returned 630 systems.

Among these systems, 9 sequences have non-standard letters (usually an X) and were removed. Another 34 structures have interruptions in the structure (such as unresolved part in the middle) and were also removed. Some non-standard base names in the structure file correspond to modifications, and we tried to guess the original unmodified based based on the ring structure. After such treatment, *N* = 577 structure-sequence pairs were left. Among these pairs, only *N*^*′′*^ = 533 of them have a sequence length *L* ≤ 512, which is the longest sequence length StructRFM can handle. However, we emphasize that REDIAL algorithm is applicable to sequences of any arbitrary length with the capability of the underlying language model.

### 2 REDIAL: RNA Embedding perturbation Diagnostics for Language models

The REDIAL framework evaluates the sensitivity of the sequence representation **z** ∈ ℝ^*L*×*c*^ to single-point mutations. A known challenge when interpreting these latent spaces is the presence of high-magnitude activations in specific channels, which can disproportionately dominate the signal^73, 74^. To mitigate this, we first perform a channel-wise normalization of the raw embeddings to ensure each feature dimension has zero mean and unit variance:

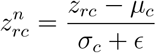

where *r* and *c* denote the sequence and embedding dimensions, respectively, and *ϵ* = 10^−12^ is a small constant added for numerical stability. This normalization is essential, as otherwise, the unbalanced activation will bury any meaningful evolutionary coupling information into noise.

For each of the *L* × *A* possible point mutations, we quantify the local response by calculating the difference in the normalized embedding, *δz*_*n*_ = *z*^*n*^(mutation) −*z*^*n*^(wild-type) . This process generates a four-dimensional sensitivity tensor 𝒥 ∈ ℝ^*L*×*A*×*L*×*c*^. To extract a site-specific interaction signal, we compute the Frobenius norm across the alphabet (*A*) and channel (*c*) dimensions, resulting in the raw interaction matrix *C* ∈ ℝ^*L*×*L*^:

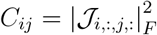

Finally, the Average Product Correction (APC) is applied to the symmetrized matrix to systematically correct for background noise and phylogenetic bias^41^:

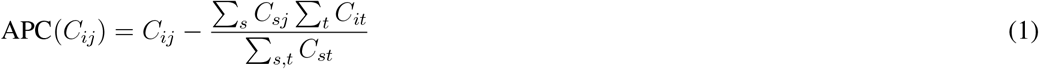

To account for the bidirectional nature of residue-residue interactions, this matrix is symmetrized by averaging it with its own transpose.

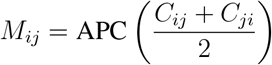

### 3 Direct Coupling Analysis

To compute DCA for the five systems we reported, we first constructed the MSA using RNACentral 24.0, which is the the same as the training set of StructRFM^55^. MSA was done with BLASTN with an E-value of 0.001.^75^ Then pyDCA was applied to compute the mean field DCA^76, 77^, using a value of sequence identity of 0.8 and a value of relative pseudocount of 0.5. APC was also applied to the results. A detailed statistics of the results were provided in Table 2. The mean field results were reported in Figure 2, and the results from pseudolikelihood maximization is in Figure S2.

**Table 2.** Statistics on MSA and DCA results.

| System | Length | MSA sequences | Effective sequences |
| --- | --- | --- | --- |
| 1FIR | 76 | 60,573 | 26.33 |
| 1Y26 | 71 | 161 | 5.84 |
| 2GDI | 80 | 1,283 | 10.68 |
| 3Q3Z | 75 | 21 | 5.75 |
| 4LVV | 85 | 58 | 4.88 |

### 4 Contact map calculation

The contact map (from 3D structures) was derived from base pairing computed with the annotation function of Barnaba^78^. A contact is marked if the annotation function suggests an interaction other than XXX, which means the two bases close but cannot be categorized into a specific type of interaction.

### 5 Evaluation of predicted coupling with PR-AUC

The challenge to evaluate the quality of the prediction is that the predicted contact does not have a fixed scale. To evaluate the accuracy of the predicted contact maps, we calculated a normalized Area Under the Precision-Recall Curve (PR-AUC).

First, we transformed the experimental distogram into a binary ground-truth contact matrix with abovementioned method. Because contact maps are symmetric, we extracted and flattened the upper triangle (excluding the diagonal) for both the ground-truth and predicted matrices to avoid redundant data. The first diagonal was also removed to avoid counting in interactions in neighboring bases. We then ranked all potential residue pairs by their predicted scores in descending order to generate a Precision-Recall (PR) curve, where recall and precision were calculated cumulatively at each rank. The Area Under the Curve (AUC) was computed from these values using the trapezoidal rule.

Finally, to account for the varied secondary structure of different systems, we calculated a normalized AUC score. For a totally random predictor, this AUC score should be equal to the ratio of true contacts to all combinations *p*. A normalization adjusts the raw AUC by subtracting this background precision

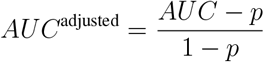

such that a score of 1 represents a perfect prediction and 0 represents a performance no better than random. Sometimes a score can be lower than 0 because of this normalization (as seen in some of Figure 5b), and that means such prediction is even worse than a random predictor.

### 6 Structural prediction of the AU-rich element with physical models

The sequence we started was GUGAUUAUUUAUUAUUUAUUUAUUAUUUAUUUAUUUAG. One previous study has employed the thermodynamic-based mFold to to identify the minimum-free-energy secondary structure, for which the majority of predicted base-pairing interactions were validated by prior experimental studies^50^. Building upon this secondary structure, the Fragment Assembly of RNA with Full-Atom Refinement (FARFAR2) algorithm is further employed^52^ to construct tertiary structures from an experimentally derived fragment library. A total of 2,000 candidate structures are generated through Monte Carlo-based refinement, and the final structural model is selected based on the lowest energy evaluated by an all-atom, knowledge-based scoring function^52^.

## Author Contributions

DT: conceptualization, methodology, data curation, software, investigation, formal analysis, writing – original draft, writing – review and editing; YQ: conceptualization, methodology, investigation, writing – original draft, writing – review and editing; GS: software, investigation, formal analysis; AA: conceptualization, methodology, writing – review and editing; LH: conceptualization, writing – review and editing; PT: conceptualization, writing – review and editing, resources, supervision, project administration, funding acquisition.

## Data Availability

The RNA language models we tested are available in their GitHub repositories: RNA-FM and RhoFold (https://github.com/ml4bio/RNA-FM) and StructRFM (https://github.com/heqin-zhu/structRFM). The code for our REDIAL algorithm is available open source at GitHub: https://github.com/tiwarylab/redial.

## Acknowledgments

P.T. is an investigator at the University of Maryland-Institute for Health Computing, which is supported by funding from Montgomery County, Maryland and The University of Maryland Strategic Partnership: MPowering the State, a formal collaboration between the University of Maryland, College Park and the University of Maryland, Baltimore. This research was supported by the National Institute of General Medical Sciences of the National Institutes of Health under Award Number R35GM142719 and the Maryland Technology Development Corporation (TEDCO) through the Maryland Innovation Initiative (MII) program under Grant No. 07250192. We thank UMD HPC’s Zaratan, Institute for Health Computing’s Beacon and NSF ACCESS (project CHE180027P) for computational resources.

## Supplementary Information

### A Comparison between Categorical Jacobian and Embedding Perturbation

“Categorical Jacobian” (CJ) algorithm is an unsupervised method proposed by Zhang *et al*.^20^ to extract coevolutionary coupling from pLMs. We adapted it to work for the RNA alphabets. The largest difference between REDIAL and CJ algorithm is that CJ looks at changes in logits, and REDIAL looks at changes in the embedding (Figure 5). The details are described as follow:

Decoders for RNA language models typically produce a logit with the dimension *L* × *A*, where the model has a vocabulary of *A* = 4 standard tokens representing the nucleotides A, U, C, and G. Similarly, because there are a total of *L* × *A* possible single-point mutations in the sequence, a four-dimensional “Jacobian”-style tensor 𝒥 ∈ ℝ^*L*×*A*×*L*×*A*^ can be defined to capture how each of these *L* × *A* potential mutations perturbs the entire *L* × *A* logits matrix. This comprehensive tensor maps every possible mutation to its effect on every possible output score.

To draw residue/nucleotide-level couplings, we first compute the Frobenius norm of the Jacobian 𝒥 across the two alphabet-specific dimensions (the letter-dimensions). This reduction yields a two-dimensional interaction map *C* ∈ ℝ^*L*×*L*^:

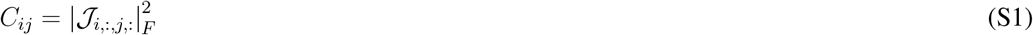

Then, the APC algorithm (Equation 1) is applied to reduce noise.

CJ algorithm is more susceptible to noise than REDIAL for RNAs. While the reduction from the 4-D tensor to 2-D coupling is robust for proteins, where the alphabet size *A* = 20 provides a high-dimensional buffer against local fluctuations, it becomes problematic for RNA (*A* = 4). The 5× smaller vocabulary means that the resulting coevolutionary matrix is more sensitive to noise in the model, leading to weakened signals.

**Figure S1.**
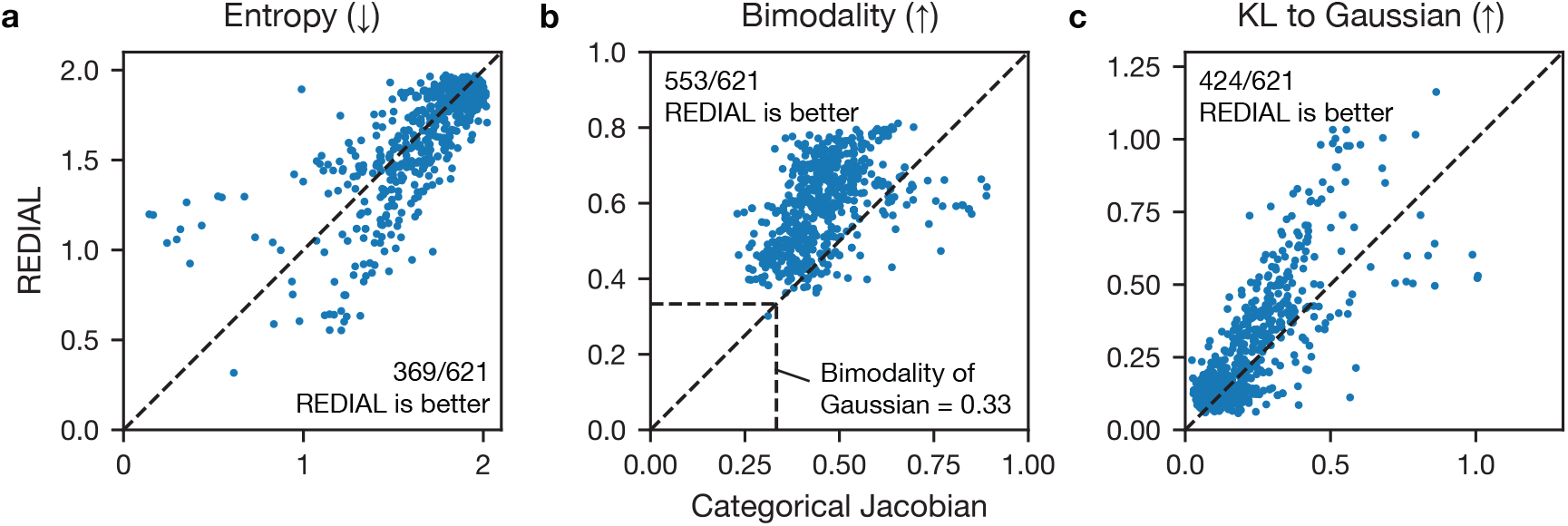
Comparison between Categorical Jacobian method and Embedding Perturbation. Three quantities were computed for the contact maps predicted from NAKB dataset we cleaned. (a) Shannon entropy, (b) Bimodality coefficient, and (c) Kullback-Leibler divergence to a Gaussian.

This higher level of noise can be quantified in several measures. We evaluates how noisy each contact map is by looking at the distributions *p*(*x*) of the values in the predicted contact *x*. First, the predicted couplings were converted to *z*-score, with zero mean and unity variance. Then, three measures were computed:

#### Shannon Entropy

The Shannon entropy defined as

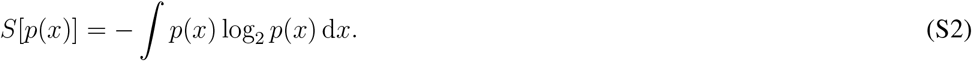

The smaller this value is, the more spread out the distribution is. In an informative contact map, the coupling signals should be distinct from the background. A lower entropy value indicates a more structured, less uniform distribution, reflecting a departure from random noise.

#### Bimodality coefficient

Sarle’s Bimodality Coefficient is defined to measure how bimodal a distribution is. The more bimodal, the more informative the predicted contact map may be. It is defined as

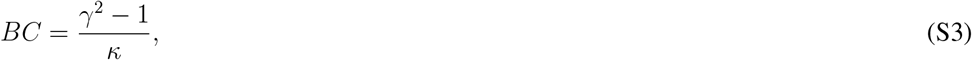

where *γ* = ⟨ (*x* − *µ*)^3^⟩ */σ*^3^ is the skewness (*µ* for mean, and *σ* for standard deviation) and *κ* = ⟨ (*x* − *µ*)^4^⟩ */σ*^4^ is the kurtosis. Specifically, a Gaussian distribution has a bimodality coefficient of 1*/*3. An ideal contact map should exhibit a bimodal distribution, where a wide “noise” peak is clearly resolved from a “signal” peak corresponding to true physical contacts. A higher bimodality coefficient thus indicates better resolution of structural features.

#### Kullback-Leibler divergence to a Gaussian

This measure how similar this distribution is to a Gaussian 𝒩(0, 1), which is different from the Shannon entropy by a cross entropy term. It is defined as

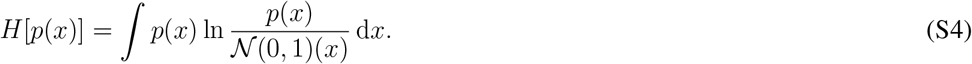

A larger KL divergence signifies that the predicted map is less likely to be a Gaussian-like stochastic noise, indicating a more robust extraction of non-random evolutionary signals.

While these metrics represent different mathematical properties of the data, in every assessment, the contact maps generated via REDIAL demonstrated statistically better performance compared to the CJ algorithm in RNAs, especially in the bimodality test. This confirms that perturbing the higher-dimensional hidden representation effectively averages out the categorical noise inherent in logit-based extractions more significant in RNA models.

### B Results of Direct Coupling Analysis (DCA) for the five systems

The result of the DCA and its multiple sequence alignment (MSA) were summarized in Supplementary Table 2. In the main text, we reported the result from mean field DCA. Here we also show show the result from Pseudolikelihood maximization DCA (Figure S2). Due to the low number of available sequences, they appear much noisier.

**Figure S2.**
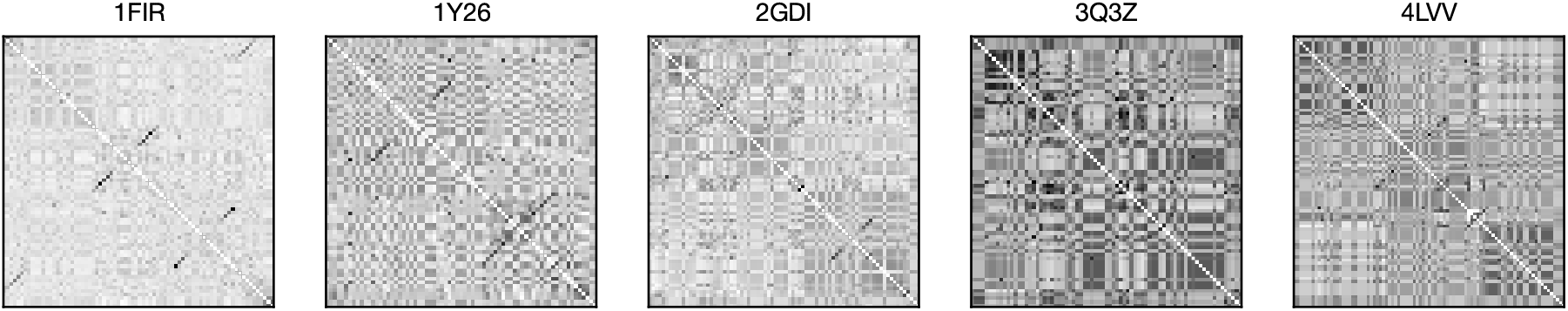
Pseudolikelihood maximization DCA results. Comparing with the meanfield DCA in the main text, these signals are noisier due to small number of available MSA sequences.

### C Statistics on Contact PR-AUC scores

The distribution of PR-AUC scores were analyzed in detail here. Figure S3 shows the total distribution shows StructRFM is much better than the vanilla RNA-FM.

**Figure S3.**
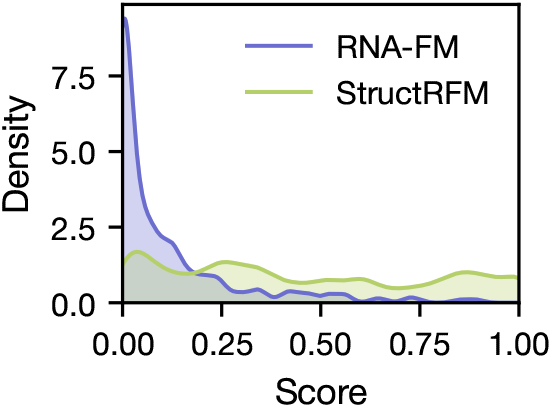
The Contact PR-AUC scores distribution for RNA-FM and StructRFM.

We dissected the scores in four categories and showed their respective performance in Figure S4. In the category of tRNAs, some shorter sequences are from anticodons of tRNAs. They are often much shorter in length.

**Figure S4.**
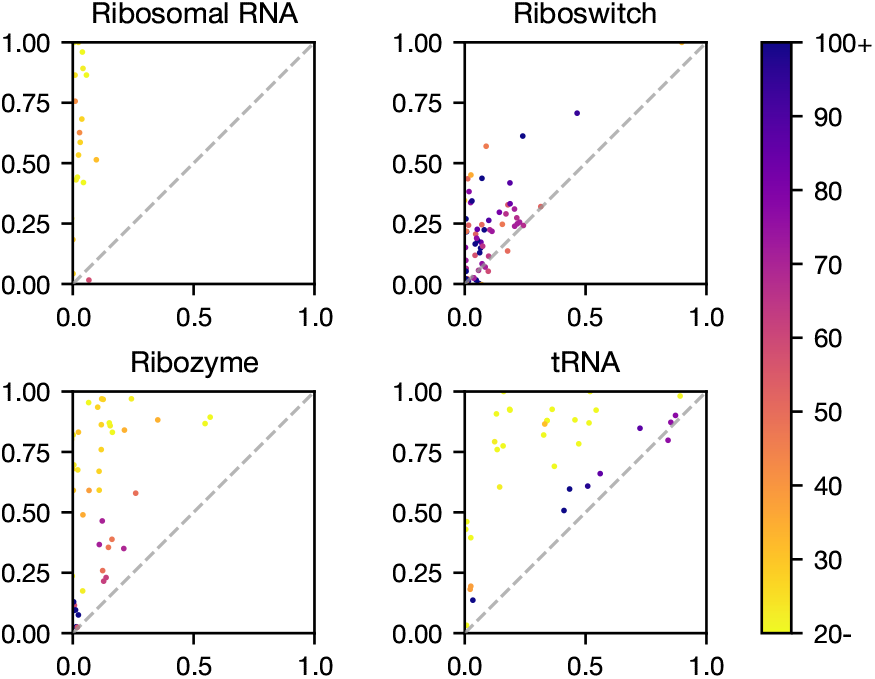
Comparison of contact scores by NAKB annotations. Note that RNAs not in these four categories are not shown here.

We also analyze the distribution of scores by length group in Figure S5. These two models show clear discrepancies in behavior regarding sequence length. StructRFM’s performance decreases with length while RNA-FM is consistently not performing.

**Figure S5.**
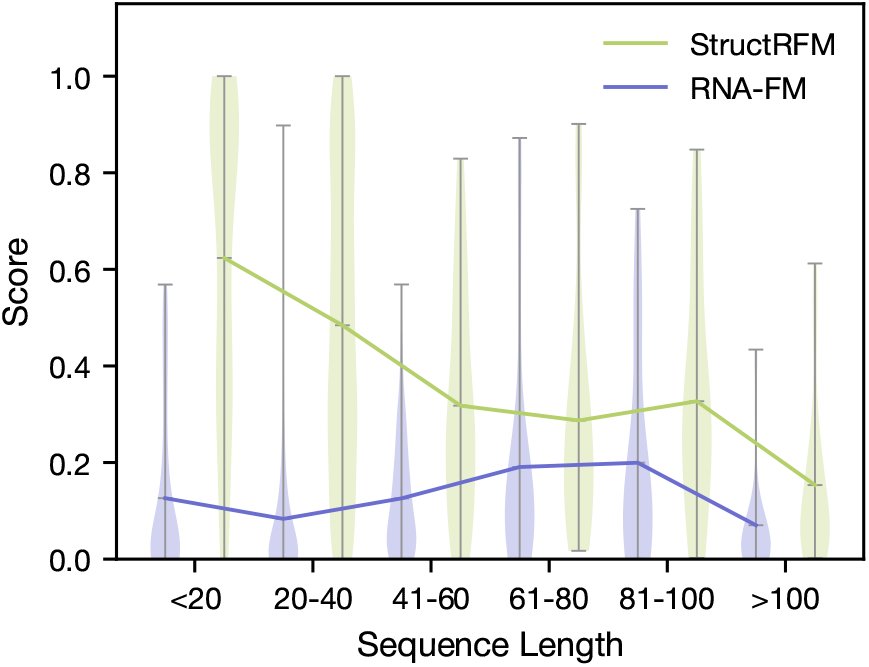
Score distribution violin plot for six length groups. StructRFM performs consistently better than RNA-FM. However, the performance of both models struggle with long RNAs with length *>* 100. StructRFM has the largest margin for short RNAs with length *<* 20.

## Additional Supplementary Figures

**Figure S6.**
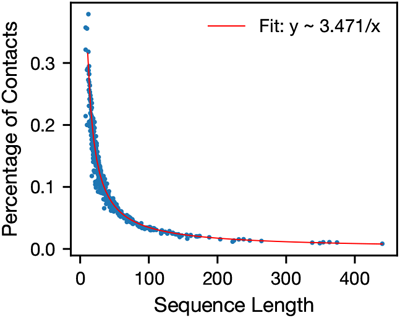
Percentage of contacts vs. sequence length. The percentage of base pairs within 6.5 Å of each other among all pairs. The fit was done with a linear fit between 1*/y* and *x*. Pearson coefficient

**Figure S7.**
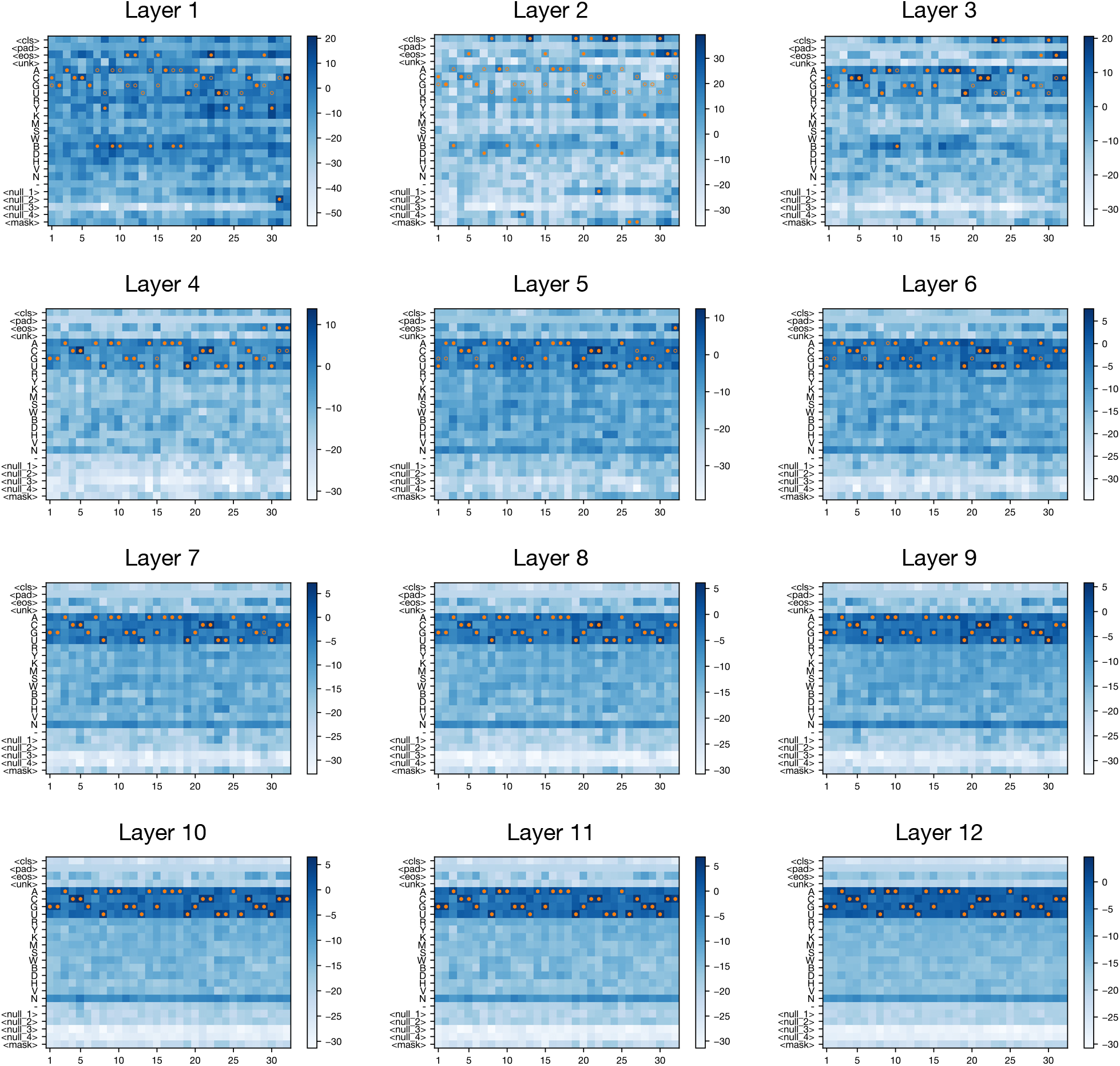
Layerwise decoded logits from 2LI4 and RNA-FM. Hollow point: ground truth sequence. Solid point: predicted token (with largest log-likelihood)

